# Molecular docking, simulation and binding free energy analysis of small molecules as *Pf*HT1 inhibitors

**DOI:** 10.1101/2022.04.27.489756

**Authors:** Afolabi J. Owoloye, Funmilayo C. Ligali, Ojochenemi A. Enejoh, Adesola Z. Musa, Oluwagbemiga Aina, Emmanuel T. Idowu, Kolapo M. Oyebola

## Abstract

Malaria chemotherapy has been plagued by parasite resistance. Novel drugs must be continually explored for malaria treatment. *Plasmodium falciparum* requires host glucose for survival and proliferation. Protein involved in hexose permeation, *P. falciparum* hexose transporter 1 (*Pf*HT1) is a potential drug target. We performed high throughput virtual screening of 21,352 small-molecule compounds against *Pf*HT1. The stability of the lead compound complexes was evaluated via molecular dynamics (MD) simulation for 100 nanoseconds. We also investigated the pharmacodynamic, pharmacokinetic and physiological characteristics of the compounds in accordance with Lipinksi rules for drug-likeness to bind and inhibit *Pf*HT1. Molecular docking and free binding energy analyses were carried out using Molecular Mechanics with Generalised Born and Surface Area (MMGBSA) solvation to determine the selectivity of the hit compounds for *Pf*HT1 over the human glucose transporter (hGLUT1) orthologue. Five important compounds were identified: Hyperoside (CID5281643); avicularin (CID5490064); sylibin (CID5213); harpagoside (CID5481542) and quercetagetin (CID5281680). The compounds formed intermolecular interaction with the binding pocket of the target via conserved amino acid residues (Val314, Gly183, Thr49, Asn52, Gly183, Ser315, Ser317, and Asn48). The MMGBSA analysis of the complexes yielded high free binding energies. Four (CID5281643, CID5490064, CID5213, and CID5481542) of the identified compounds were found to be stable within the *Pf*HT1 binding pocket throughout the 100 nanoseconds simulation run time. The four compounds demonstrated higher affinity for *Pf*HT1 than the human major glucose transporter (hGLUT1). This investigation demonstrates the inhibition potential of sylibin, hyperoside, harpagoside, and avicularin against *Pf*HT1 receptor. Robust preclinical investigations are required to validate the chemotherapeutic properties of the identified compounds.

## Introduction

Malaria is a significant cause of mortality and morbidity in Africa. A 2021 report estimated about 229 million cases of malaria and approximately 409, 000 deaths worldwide [1]. Malaria burden remains a global concern despite concerted efforts to mitigate the transmission of the parasites. Although different classes of antimalarial drugs such as quinine, antifolates, artemisinin and its derivatives have been used to treat malaria with great success, the emergence of multi-drug-resistant *P falciparum* strains has necessitated the development of new therapeutic options [2–4].

There is a need to develop novel antimalarial drugs with high potency, favourable pharmacodynamic and pharmacokinetic profiles, effective solubility, metabolic stability, permeability and transporter effects. With recent advances on the molecular mechanisms of parasite resistance, new development opportunities have emerged for the design and implementation of effective mitigation strategies [5, 6]. Protein-based rational drug design relies on high-resolution target structures to allow for high-throughput screening of selective ligands either to agonize or antagonize such target structures [7]. Proteins such as *P falciparum* hexose transporter (*Pf*HT1) and *P falciparum* dihydroorotate dehydrogenase (*Pf*DHODH) are essential for parasite survival and proliferation [8, 9]. The malaria parasites depend solely on the host’s glucose for the primary carbon source of biomass production and ATP synthesis [10].

A useful strategy for the development of new antimalarial drugs involves selectively antagonizing the parasitic carbon source without compromising the host cell’s physiological integrity [11]. *Pf*HT1 scavenges sugar from an infected host’s erythrocytes to enhance the parasite’s chances of survival and proliferation [12]. The parasite has a significant survival advantage due to the discovery of the *Pf*HT1 binding site, which facilitates the protein’s capacity to transport a wide spectrum of sugar molecules successfully [9]. *P. falciparum-infected* erythrocytes utilize up to 100 times more glucose than non-infected erythrocytes require because the parasite continuously metabolises sugars from the host’s erythrocytes to support its survival, growth, and replication [13].

In this study, we performed structure-based high throughput screening of 21,352 phyto-ligands libraries designed specifically to follow Lipinski’s rules for potential small drug molecules [14]. We also performed molecular dynamics simulations on the lead compounds to elucidate protein motion by following their conformational changes through time. In all, we identified bioactive hit-to-lead compounds with varying potency and selectivity for *Pf*HT1 over the human glucose transporter (hGLUT1).

## Materials and methods

### Protein Preparation

The 3-dimensional (3D) structure of *Plasmodium falciparum* Hexose Transporter 1 protein (*Pf*HT1) was retrieved from Protein Data Bank (PDB) (https://www.rcsb.org). The *Pf*HT1 protein with PDB ID 6M02 was refined by using Protein Preparation Wizard of Schrödinger-Maestro Release 2021-4 [15]. We assigned charges, bond orders, and deleted water molecules. Hydrogens were added to the heavy atoms. We performed energy minimization using Optimized Potentials for Liquid Simulations (OPLS) 2005 force field by fixing the heavy atom root-mean-square deviation (RMSD) of 0.30Å [16]. Lastly, we optimized amino acids using neutral pH.

### Ligand Preparation

A total of 21,352 ligands from plants that have been documented to have antiplasmodial activities were used. The phytochemicals were downloaded in the structure-data file (SDF) format from the NCBI PubChem database (https://pubchem.ncbi.nlm.nih.gov/). Ligand preparation was performed on the downloaded SDF files to assign proper bond orders and create a three-dimensional geometry [17]. This was done in Maestro Schrodinger Suite 2017 using the Ligprep with OPLS 2005 force field [17]. Ionization states were generated at pH 7.0 ± 2.0 with Epik 2.2 in Maestro Schrodinger Suite 2017 [18]. We generated 15 possible stereoisomers per ligand.

### Receptor Grid Generation

The receptor grid was generated on the prepared protein. OPLS 2005 force field was used to generate the grids [19]. The van der Waal radii of the protein atoms were scaled by 1.0, the charge cutoff for polarity was 0.25. The receptor grid box was generated in each direction (x = 27Å, y = 27Å, and z = 27Å). The dock main after the grid generation was 72Å for each dimension (x, y and z).

### Standard Precision (SP) and Extra Precision (XP) Ligand Docking

The molecular docking experiment was performed employing the Glide as executed in the Schrödinger suites [16], the receptor is treated as a stiff structure, but the ligands are treated as flexible. The receptor grid was given a dimension suitable to accommodate ligand structures with a length ≤ 14Å and a cubing docking grid was centered on Val318. The van der Waals scaling factor was set to 0.85 and 0.15 for non-polar atoms of the ligand and the partial charges limit value was set at −10.0 kcal/mole. High Throughput Virtual Screening (HTVS) and Standard Precision (SP) scoring functions of glide were used, and ligands were granted full flexibility. A post-docking minimization was carried out on output ligand-receptor complexes, reducing the initially collected 15 poses per ligand to 5. The SP resultant compounds were further docked using Extra Precision (XP) mode with more accuracy and computational intensity. The configuration was without minimization, relaxation, or flexibility. Based on the glide energy and XP glide rescoring, the procedure gives the lead ligand-receptor complexes. The glide module of the XP visualizer interface was used to examine the specific interaction. Interaction energies between ligands and proteins, hydrogen bonds, hydrophobic interactions, internal energy, pi-pi (π-π) stacking interactions, and RMSD were all included.

### Prime Molecular Mechanics with Generalised Born and Surface Area Solvation (MM-GBSA)

Binding free energy calculation was performed for the ligand-receptor complexes using the Schrodinger suite MM-GBSA module integrated with Prime [16]. The binding free energy of XP Glide docked output complexes were evaluated using Prime MMGBSA. The evaluation of the complexes’ relative energy was done with the OPLS3 force field and rotamer algorithm [20]. The free binding energy equation is: ΔGbind = Gcomplex – X – (Gprotein + Gligand). A more negative score signifies a stronger binding.

### Molecular Dynamics Simulation

Molecular dynamics (MD) simulations were performed for 100 nanoseconds using Desmond Schrödinger [16, 21]. Protein-ligand complexes used for molecular dynamics simulation were obtained from docking studies to provide a prediction of ligand binding status in static conditions. Since docking is a static view of the binding pose of a molecule in the active site of the protein, MD simulation tends to compute the atom movements with time by integrating Newton’s classical equation of motion [22]. The ligand binding status in the physiological environment was predicted using molecular dynamics simulations. Protein Preparation Wizard of Maestro Schrodinger Suite 2017 was used to preprocess the protein–ligand complex, which comprised complicated optimization and minimization. All systems were prepared by the System Builder tool. The OPLS_2005 force field was used in the simulation. Solvent Model with an orthorhombic box was selected as Transferable Intermolecular Interaction Potential 3 Points (TIP3P). We neutralized the models by adding counter ions where necessary. To mimic the natural physiological conditions, 0.15 M NaCl was added. The NpT ensemble with 300 K temperature and one atmospheric pressure was selected for complete simulation via Martyna–Tuckerman–Klein barostat [23]. The models were relaxed before the simulation and the trajectories were saved after every 100 ns for analysis, after which the stability of simulations was evaluated by calculating the RMSD of the protein and ligand over time. RMSF and protein-ligand contacts were also analyzed.

### ADME-Tox Properties

For the analysis of the pharmaceutical, physiological, biochemical, and molecular effects of the compounds, adsorption, distribution, metabolism, excretion, and toxicity (ADME-Tox) properties were calculated with the QikProp program of Maestro Schrodinger Suites [24]. The QikProp predicted the physicochemical and pharmacokinetic properties of the compounds. It also assessed the tolerability of the analogues based on Lipinski’s rule of five (that is, it does not violate more than one of the following criteria; no more than five hydrogen bond donors; no more than ten hydrogen bond acceptors; a molecular mass of less than 500 daltons; and a log *P* of less than five for octanol-water partition coefficient) [25].

### Data Analysis

The raw trajectory files from the MD simulation run time were analyzed in R (version 4.0.5) and visualized using “ggplot2” and “ggrepel” packages.

## Results

### Molecular Docking, MMGBSA/Prime Binding Energy of *Pf*HT1-Ligand Complexes

We screened a library of 21,352 compounds against the target protein, *Pf*HT1, and identified five hits from 437 compounds (S1 File) with excellent docking scores after thorough validation using the Lipinski rule of five. While *Pf*HT1 and the human major glucose transporter, hGLUT1 share some sugar transporter features, important mechanistic differences in their interaction with substrates were identified. *Pf*HT1 can transport both D-glucose and D-fructose whereas hGLUT1 is selective for D-glucose. To prove our hit compounds selected for *Pf*HT1 over hGLUT1, we docked the five hit compounds into the binding pocket of hGLUT1 and found that hGLUT1 has a very low affinity for the substrates. This implies that the compounds only bind successfully to the target protein. The molecular docking study of the five selected compounds and the target revealed that hyperoside had the highest glide docking score (−13.881). The other four ligands; sylibin, avicularin, quercetagetin and harpagoside had −12.254. −11.952, −11.756 and −11.258 docking scores respectively (Fig 1A). For the residue interactions of a protein molecule with the ligand compounds, we analyzed the protein-ligand complex structure and discovered that Val314 formed a hydrogen bond with four of the five complexes, while Ser317 and Gly183 appeared in three complexes with a hydrogen bond. Ser315, Asn48, Asn52, and Thr49 formed hydrogen bonds with two complexes each Table 1. This suggests that the amino acid residues are essential for target binding.

**Fig 1.**
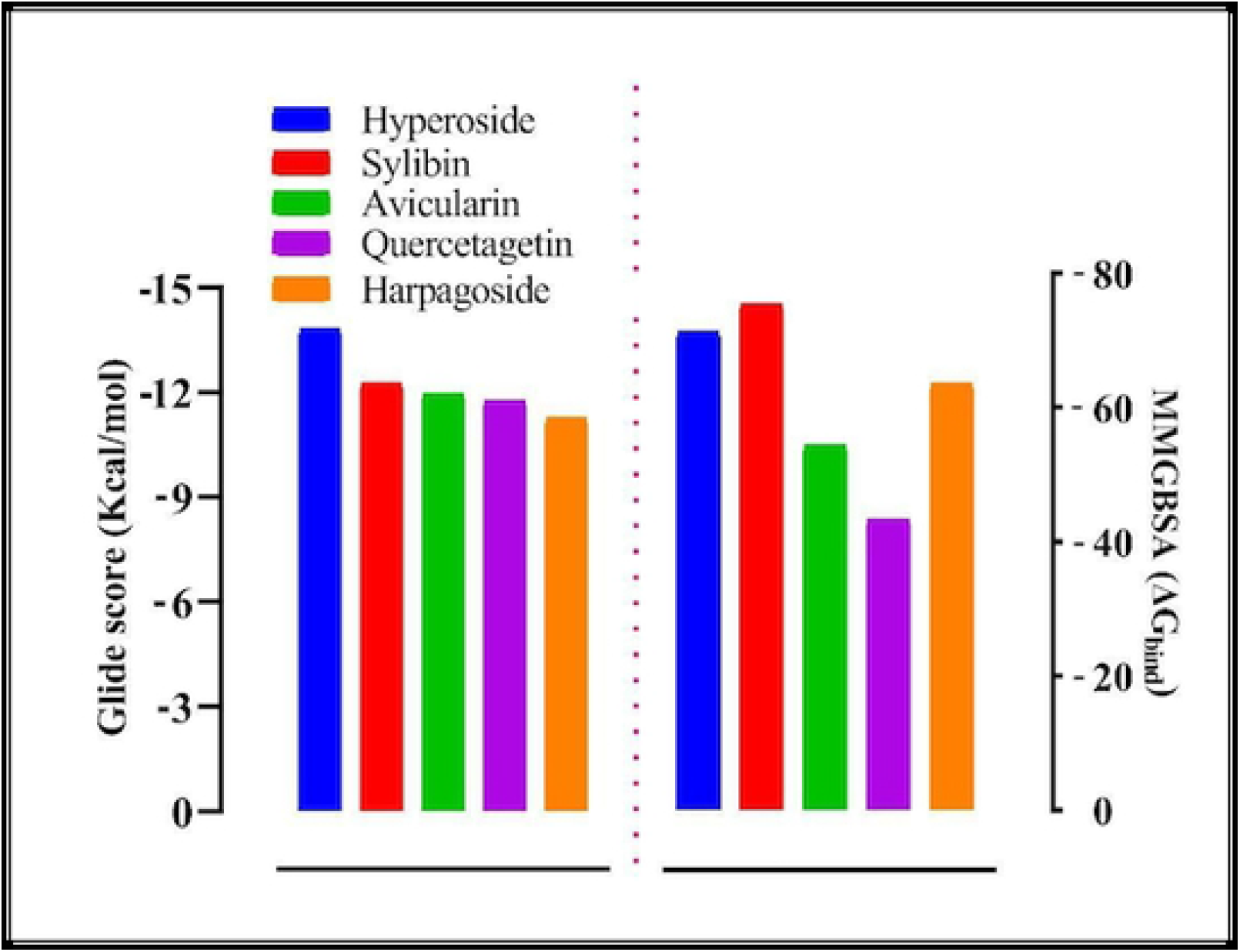
(A) Molecular docking glide score (Gscore) and (B) Prime MMGBSA binding energy (ΔG_bind_) of the lead compounds with PfHT1 (6m20).

**Table 1:** Intermolecular interaction of protein-ligand complexes following molecular docking.

| Compound | Hbond (distance) | Pi-pi (distance) | Arom H-bond (distance) |
| --- | --- | --- | --- |
| Sylibin | Val314 (1.7), Gly183 (2.02), Thr49 (2.11) | Asn52 (1.01), Ser317 (0.96) | Phe53 (2.30), Asn48 (2.35) |
| Hyperoside | Asn52 (2.19), Gly183(2.14), Val314(1.57; 2.13), Ser315 (0.96), Ser317 (1.52) | Asn318 (1.42) |  |
| Avicularin | Asn48 (2.02), Ser315(2.60), Ser317 (1.67), Val314 (2.57; 1.63), | Asn48 (1.95) |  |
| Quercetagenin | Val314 (1.5; 2.26) | Asn48 (2.88), Asn318 (3.00), Ser317 (1.85) |  |
| Harpagoside | Gly183 (1.90), Thr49 (2.18), Asn52 (2.62; 1.82), Asn48 (2.18), Ser317 (1.82) | Asn48 (2.18), Phe53 (1.74), Asn318(1.59; 2.12; 2.14) | Val314 (2.27) |

### hGLUT1 and Ligands

To determine the selectivity of our compounds as *Pf*HT1 inhibitors versus, we downloaded the hGLUT1 (6THA) protein from the protein data bank https://www.rcsb.org, prepared the protein using the protein preparation wizard of Schrodinger suite. Subsequently, a receptor glide grid was generated and molecular docking of the receptor with our hit compounds was performed. We discovered that the glide docking scores of the ligand-receptor of the five compounds were relatively low. The docking scores were −6.324, −6.065, −5.812, −5.728 and −5.133 compared to −11.952, −11.258, −13.881, −11.756, and −12.254, respectively for avicularin, harpagoside, hyperoside, quercetagetin and sylibin. Interestingly, the amino acid residues involved in the intermolecular interaction of the receptor-binding pocket were not consistent unlike *Pf*HT1. When we performed MMGBSA/Prime on the complexes to evaluate the binding free energy, we observed that the binding free energy of the complexes was also relatively low (Fig 2).

**Fig 2.**
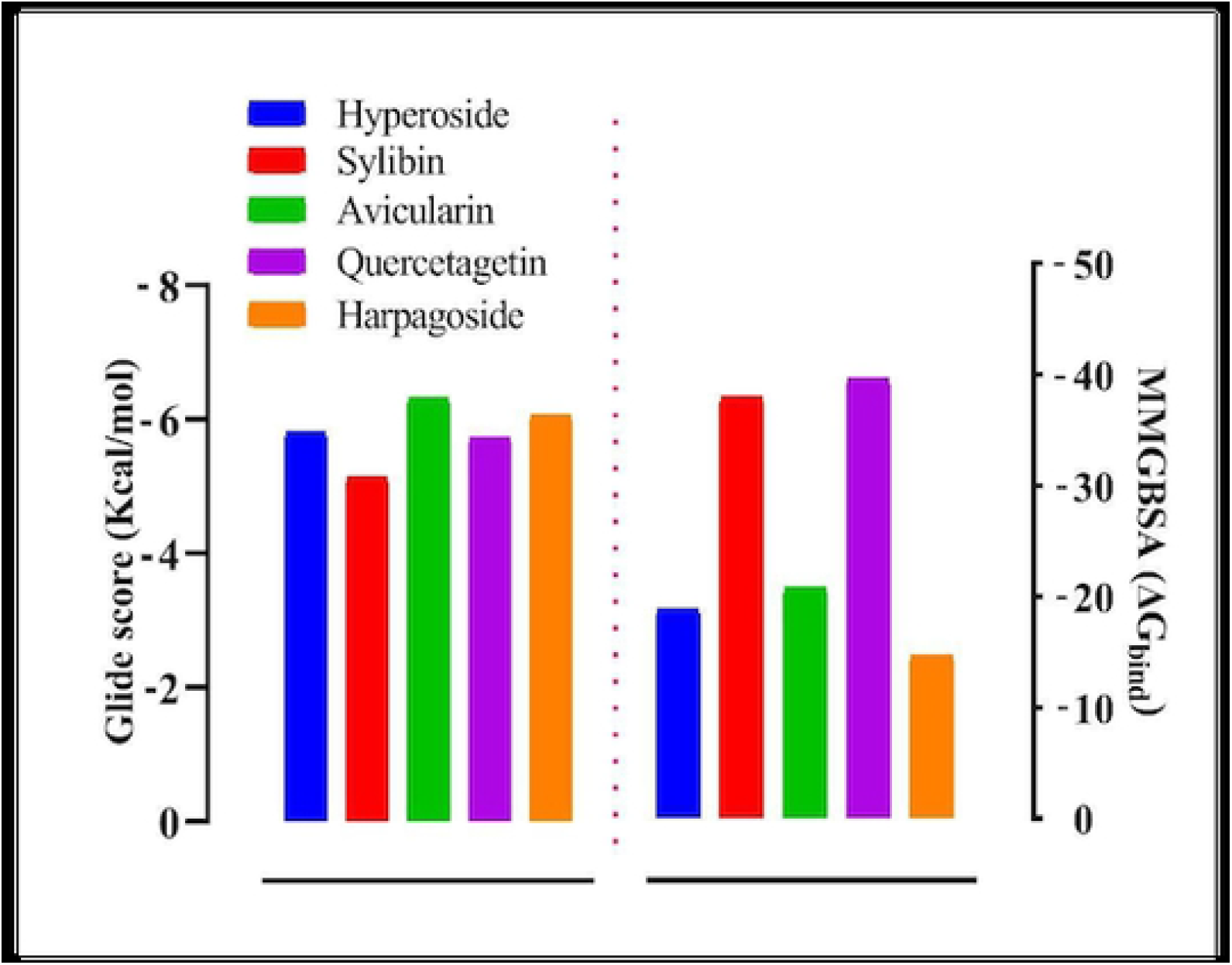
(A) Molecular docking glide score (Gscore) and (B) Prime/MMGBSA binding energy (ΔG_bind_) of the lead and with hGLUT1 (6THA).

### Interaction of Avicularin with *Pf*HT1

Among the five hit compounds, avicularin showed a glide docking score that was closest to sylibin at - 11.952. When the protein-ligand complex and the ligand atoms’ contact with the target residues were observed, we found that the residue interactions of Ser315, Ser317, Asn48, and Val314 had double H-bonds. The interaction involved back backbone and side-chain contacts, as well as hydrophobic contacts (Fig 3A).

**Fig 3.**
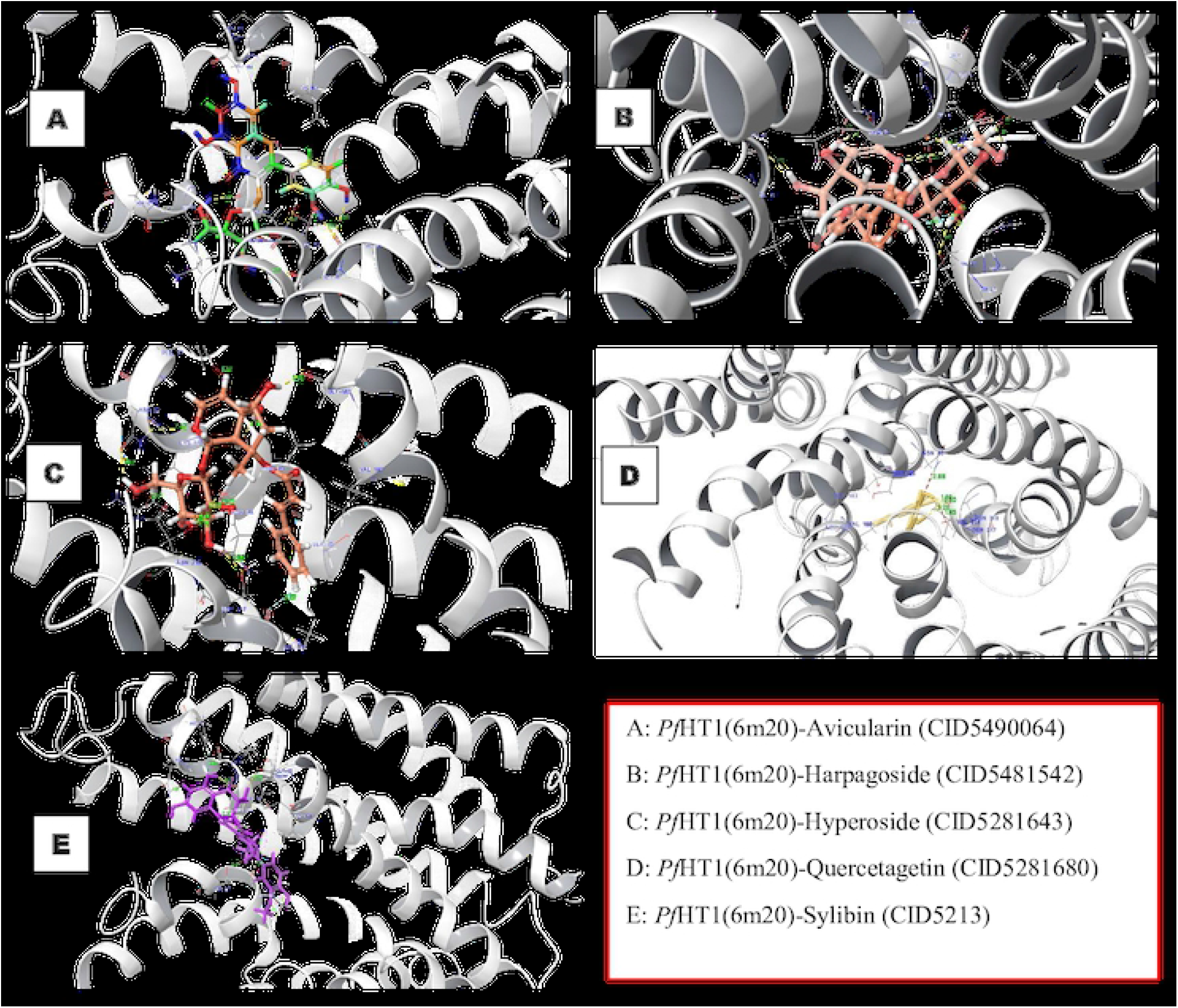
The 3D structures of interaction profile of PfHT1(6M20)–ligand complexes after molecular docking studies.

### Interaction of Harpagoside with *Pf*HT1

The *Pf*HT1 target residues interacted with the atoms of the compound, the binding surface was controlled by a range of intermolecular interactions. The binding affinity depends on interactions at the bindings site and the non-specific forces outside the target binding region. The pattern of interaction between *Pf*HT1 and harpagoside in the complex is shown in Fig 3B. The amino acid residue, Val314 formed triple pi-pi interaction with the ligand. We examined the interaction of harpagoside within the binding pocket of the target and discovered that the interaction was vigorous unlike hGLUT1 (Fig 4). This could be a result of the number of intermolecular interactions and the distance of the bonds. *Pf*HT1 residues bound to the ligand through Asn48, Try49, Gly183, Ser317, and Asn52 made double H-bonds. Asn52 formed H-bond back chain contacts with the harpagoside molecule (Fig 3B).

**Fig 4.**
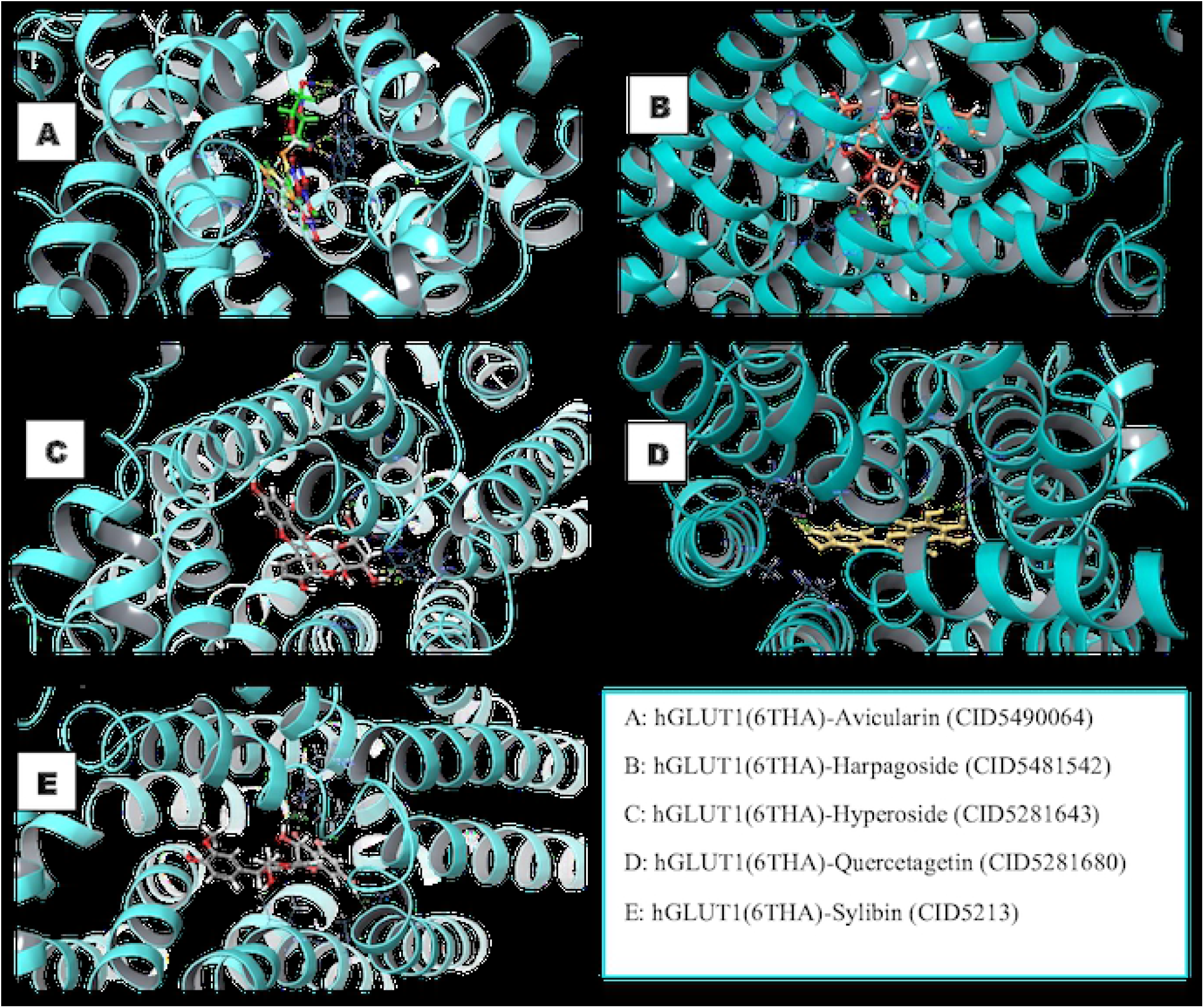
The 3D structures of interaction profile of hGLUT1(6THA)–ligand complexes after molecular docking studies.

### Interaction of Hyperoside with *Pf*HT1

The interaction of hyperoside with *Pf*HT1 is shown in Figure 1B. The docking score of the *Pf*HT1-hyperoside complex was −13.881 kcal/mol. The complex formed six conventional hydrogen bonds with Ser315, Ser317, Asn52, Gly183, and Val314 made a double hydrogen bond with the ligand (Fig 3C). The distance, 3 Å, encircled around the ligand was considered for screening. A pi-pi bond was also observed in Asn318 residue.

### Interaction of Quercetagetin with *Pf*HT1

Quercetagetin had the fourth-highest glide docking score of −11.756 and a good glide energy value. However, it is the only ligand with the least target residue contact. The target-ligand complex interaction template showed it had double H-bond residue interactions with Val314. We also observed pi-pi interaction via Asn318 and Asn48 amino acid residue. Meanwhile, the only pi-stack bond was on Ser317 (Fig 3D).

### Interaction of Sylibin with *Pf*HT1

Compound Sylibin (CID5213) occupied the binding pocket of *Pf*HT1 with the second-highest glide docking score of −12.254 Kcal/mol. Three hydrogen bond interactions were identified with the backbone amino acid residue Thr49, Gly183, and Val314 (Fig 3E). Gly183 and Val314 form H-bond contact with the backbone, while Thr49 forms an H-bond with the side chain. The interaction of the protein-ligand complex was robust as the ligand fits in perfectly into the binding pocket of the target.

### Prime MMGBSA

The prime MMGBSA integrated with the Prime Schrodinger suite was used to compute the binding free energy of the docked complexes. The relative free binding energy of sylibin, hyperoside, harpagoside, avicularin, and quercetagetin were −75.43, −71.32, −63.62, −54.41, and −24.31, respectively, as shown in Fig 1. The free binding energy further established the binding affinity of the selected ligands compared with the reference compound.

### Molecular Dynamics Simulation

The selected ligands were evaluated for their conformational stability within the receptor’s binding pocket. We examined the protein-ligand root mean square deviation (RMSD), protein root means square fluctuation (RMSF), ligand RMSF, protein secondary structure, protein-ligand contacts, and ligand torsion profile. The RMSD is the average deviation in the displacement of a group of atoms in relation to a reference frame for a given frame. Avicularin-receptor complex (lig-fit-prot) reached equilibrium after the first 30ns, the equilibrium was maintained till the end of the evolution with minimum and maximum values of 1.39 and 2.77Å, respectively. The Cα atoms reached a consistent fluctuation only after 0.5Å and was maintained all through the model (Fig 5). Likewise, the Cα atoms of sylibin, hyperoside, and harpagoside complex maintained an equilibrium state throughout the evolution time. There was a fair equilibrium in Cα atoms of the quercetagetin complex. Sylibin lig-fit-prot was observed to maintain a stable equilibrium for the period of 98 ns (Fig 5E). While hyperoside and harposide lig-fit-prot reached equilibrium after 30 ns, the steady-state was maintained for 70ns. Quercetagetin complex, on other hand, showed a steady state from 15ns to 78ns, the oscillation was between 1.6 Å and 3.2 Å. We further observed a slight fluctuation between 76 and 90ns (Fig 5D). Meanwhile, the atoms jumped back to their original state after 90ns and were maintained to 100ns. The Root Mean Square Fluctuation (RMSF) characterizes local changes along the protein chain. The *Pf*HT1 amino acid residue local changes were monitored for 100 ns simulation run time. The maximum loop region fluctuation recorded was 5.4 Å in all the models (S1 Fig). Interestingly, there was no substantial fluctuation in the loop regions while comparing within models. The amino acid residues in avicularin, hyperoside, sylibin, and harpagoside model oscillated with little fluctuations; 0.5 - 1.5 Å, 0.5 - 1.6 Å, 0.5 - 1.0 Å, and 0.6 – 1.2 Å, respectively.

**Fig 5.**
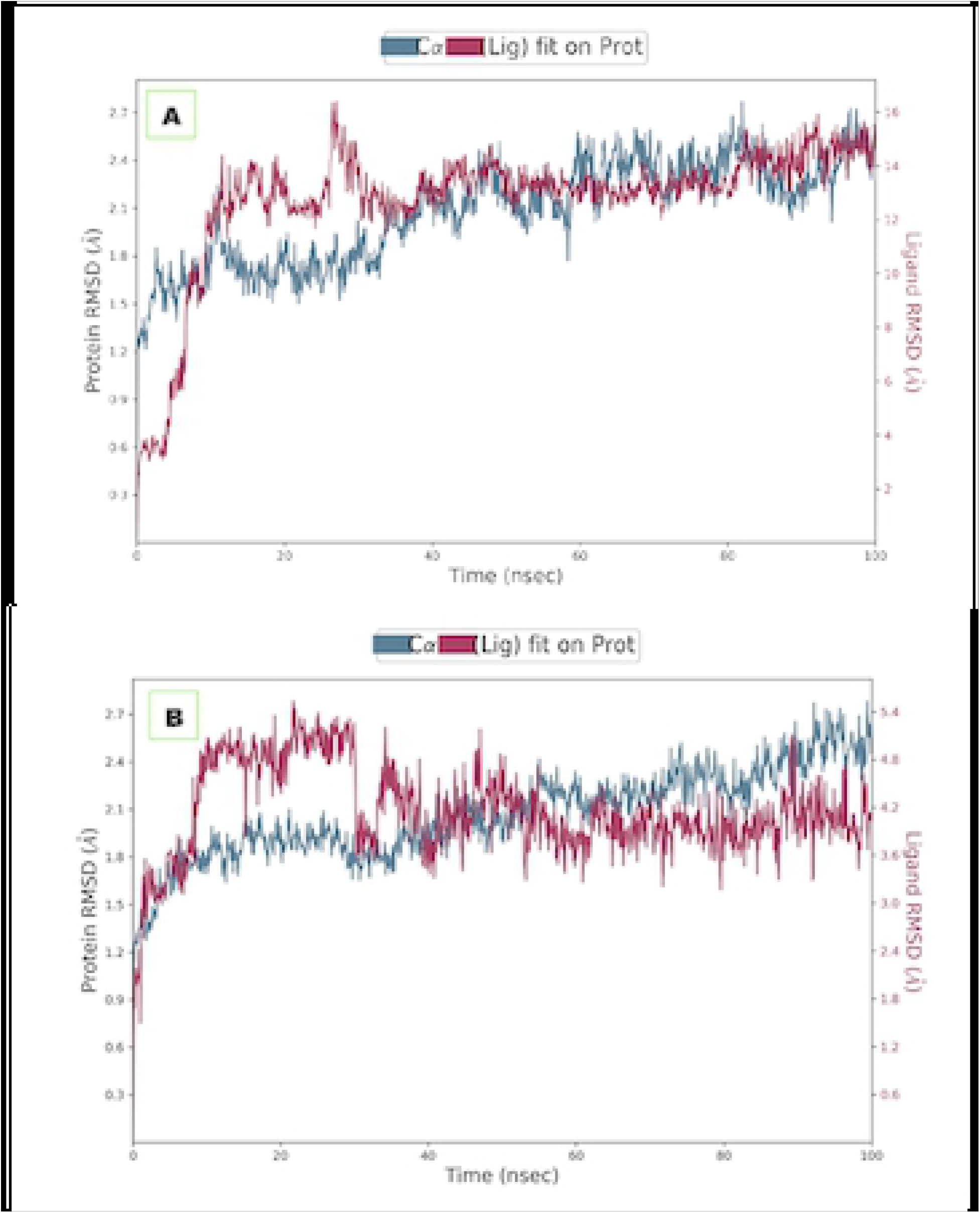
Line representation of the evolution of RMSD throughout the MD simulations of the *Pf*HT1(6m20) complex with the lead compounds. (**A**: Avicularin CID5490064; **B**: Harpagoside CID5481542; **C**: Hyperoside CID5281643; **D**: Quercetagetin CID5281680; **E**: Sylibin CID5213). The left frames show RMSD value for *Pf*HT1 - Cα, whereas the right frame shows the ligand RMSD value. Lig fit Lig illustrates the RMSD of the ligand that is aligned and measured on its reference (first) conformation

Protein-ligand interactions were monitored throughout the simulation (Fig 6 and S2 Fig). Hydrogen bonds play an essential role in protein-ligand binding. The conserved residues that formed hydrogen bond interactions were Asn48, Ser315, Lys51, Asn316, and Val 444. Interestingly, none of these residues interacted with quercetagetin, as shown in the RMSD and RMSF. Instead, quercetagetin showed remarkable RMSD and RSMF instability. Notable residues that form hydrophobic interaction with the ligands were Leu75, Leu81, Val443, and Val444, while more than 18 amino acid residues were conserved via water-bridge interactions with the ligands.

**Fig 6.**
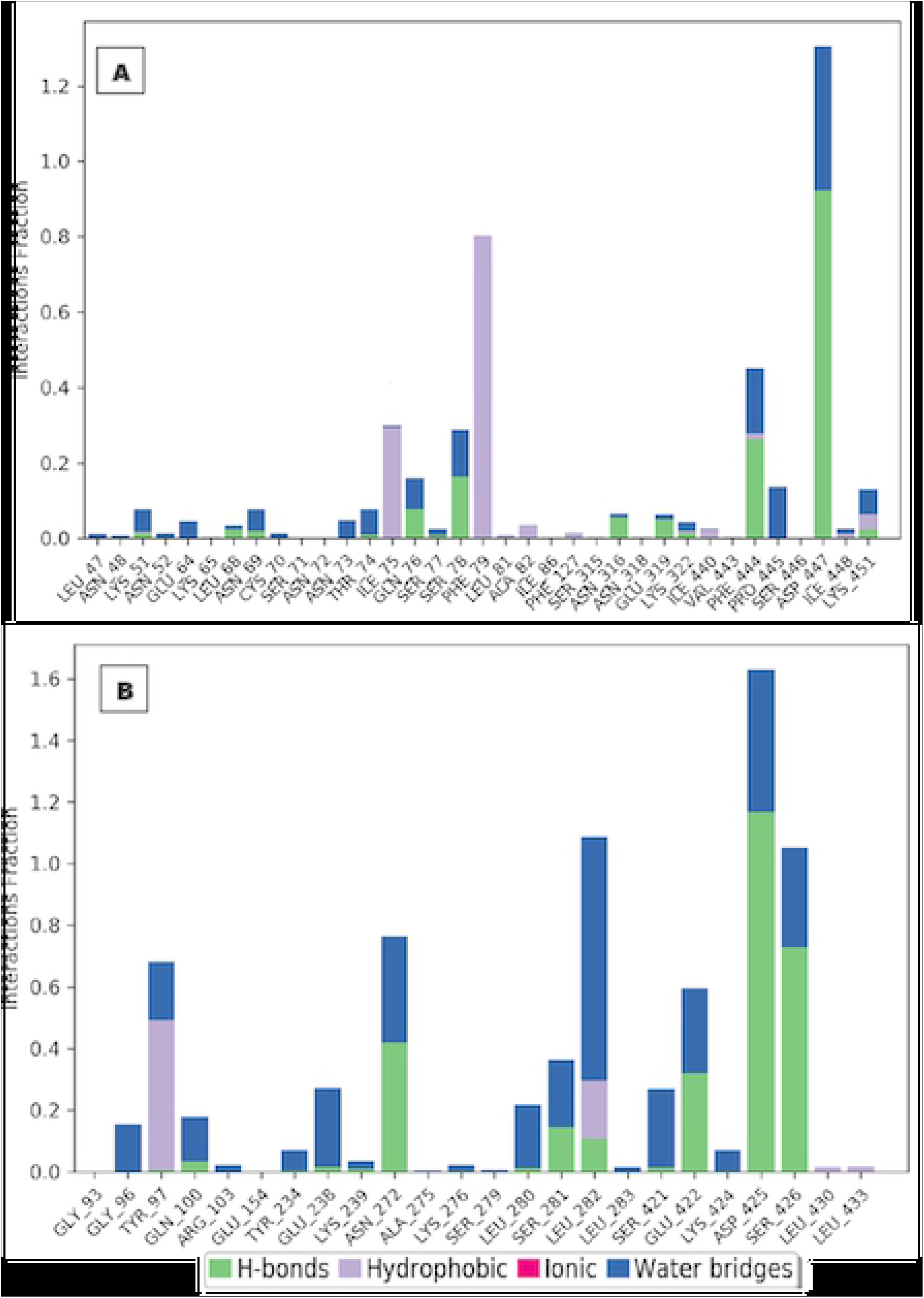
Percentage protein-ligands contacts monitored throughout 100 ns simulation run time of *Pf*HT1(6m20) complex with the lead compounds. (**A**: Avicularin CID5490064; **B**: Harpagoside CID5481542; **C**: Hyperoside CID5281643; **D**: Quercetagetin CID5281680; **E**: Sylibin CID5213) The stacked bar charts show the different types of interactions between the ligands and the *Pf*HT1 complex. The legend above depicts the type of interactions that occurred by means of the colour code.

### ADME-Tox Evaluation

We evaluated the pharmacological and pharmacokinetic features of the hit compounds to predict their physiochemical characteristics. The properties represent the absorption, distribution, metabolism, and excretion (ADME) of the compounds. To estimate the ADME properties, we employed the Lipinski rule five (RoF: molecular weight (MW < 500); hydrogen bond acceptor (HBA < 10); hydrogen bond donor (HBD < 5); and predicted octanol/water partition coefficient (QPlogPo/w<5)) [25]. As shown in Table 2, sylibin, hyperoside and harpagoside did not violate any of the Lipinski rules. This makes the compound potentially druggable. About 75%, 62%, and 58% of hyperoside, sylibin, and harpagoside, respectively will be optimally absorbed into the system. The human oral absorption (HOA) of the three compounds showed cell permeability with considerable efficiency. Although avicularin had the highest HOA (79%), it violated one rule (hydrogen donor = 5). On the other hand, quercetagetin violated two rules (accptHB > 10 and donorHB > 5), and its HOA is very low compared with other compounds. This potentially indicates that the body can only absorb 10% of the quercetagetin.

**Table 2:** ADME properties of the compounds.

| Compound | MolW | donorHB | accptHB | QPlogPo/w | HOA (%) | RoF |
| --- | --- | --- | --- | --- | --- | --- |
| Hyperoside | 464.382 | 4 | 9 | -1.126 | 75.13 | 0 |
| silybin | 482.443 | 4 | 9 | 1.628 | 62.782 | 0 |
| avicularin | 434.356 | 5 | 10 | -0.797 | 79.097 | 1 |
| quercetagenin | 318.239 | 6 | 11 | -0.259 | 10.304 | 2 |
| Harpagoside | 494.494 | 4 | 8 | -0.454 | 58.365 | 0 |

**MolW**, molecular weight (< 500KDa), **donorHB**: hydrogen donor (< 10), **accptHB**: hydrogen acceptor (< 10), **QPlogPo/w**: octanol-water partition coefficient (< 5), **HOA**: Human Oral Absorption (< 50%), **RoF**: rules of five (number of violations of Lipinski's rule of five)

## Discussion

The obstruction of the glucose uptake pathway to starve out malaria parasites serves as a strategic way of antimalarial drug discovery. *Pf*HT1 plays an essential role in the survival, proliferation, and other metabolic activities of the parasite. In this computational study, we screened a library of 21,352 compounds against *Pf*HT1 and identified five phyto-ligand inhibitors of the protein. Interestingly, the five drug-like molecules identified have been previously reported to be active against some other ailments. For example, hyperoside (CID5281643), a quercetin3-O-D-galactoside (*i.e.*, quercetin with a beta-D-galactosyl residue attached at position 3), is a flavonol glycoside present in a variety of vegetables and fruits [26]. It is predominant in *Hypericum mysorense* [27]. Hyperoside has been reported to have neuroprotective effects [28], cardio-protective activity [29] and antioxidant activity [30, 31]. Avicularin (CID5490064) or quercetin glycoside, is a plant flavonoid, (Fujimori and Shibano, 2013). It is isolated predominantly from *Foeniculum vulgare* and *Juglans regia* [32]. Avicularin, as a plant metabolite, has a hepatoprotective property [32] and has been demonstrated to reduce C/EBP-activated GLUT4-mediated glucose uptake in adipocytes, inhibiting the formation of intracellular lipids [33]. Harpagoside (CID5481542), on other hand, is a terpene glycoside chiefly from *Harpagophytum procumbens* (devil’s claw). It has been reported that the compound has anti-inflammatory properties [34] against knee osteoarthritis [35]. Quercetagetin (CID5281680), a hexahydroxyflavone, is a plant metabolite that is predominant in marigold *(Tagetes erecta)* and *Neurolaena lobata.* It has a role as an antiviral [36]; an antioxidant agent [37] and *in vitro* antilipemic potentials [37].

All five hits had similar interaction with the target compounds and fit into the target binding pocket with similar conformations, glide docking scores and very similar binding energies. The ligands had pi-pi interactions with Asn52, Asn318, Asn48, Ser317, and Phe53 residues and aromatic H-bond with Phe53, Asn48, and Val314 residues. Multiple H-bonds were present in all the docked complexes with residues Val314, Gly183, Thr49, Asn52, Gly183, Ser315, Ser317, and Asn48. Residues Val314, Ser317, and Gly183 formed an H-bond with 4 of the 5 ligand-receptor complexes. Likewise, Asn48, Asn52, and Thr49 residues had H-bond intermolecular interaction with 3 of the 5 ligand-receptor complexes. These residues play important roles as they interact with ligands as hydrogen donor, hydrogen acceptor, and pi-pi interaction. Fonseca *et al.* [38] reported similar amino acids as essential residues in the binding pocket of *Pf*HT1. The pi-pi interactions in most of the complexes are formed by Asn48, Asn52, Asn318, and Ser317. This observation is essential for drug development, as most of the H-bond residues served as both donors and acceptors. The intermolecular features observed in this study could be explored to optimize ligand-receptor complexes. This can be utilized in the synthesis of entirely new molecules capable of interacting and inhibiting the target, with biological activity *in vitro* and *in vivo* [39].

MMGBSA analyzed the ligand-receptor intermolecular interactions by determining the ligand-receptor energy values and intermolecular interactions. With a high binding score and similar binding energy, the five compounds can bind *Pf*HT1 receptors [40]. Our results have demonstrated a statistical correlation to experimental binding affinity when compared with extra precision glide docking score [41]. Taking together, the MMGBSA binding affinity, intermolecular pi–pi, and H-bond interactions with the conserved amino acid residues of *Pf*HT1, sylibin, hyperoside, harpagoside, and avicularin are potential inhibitors.

To evaluate the stability of the protein-ligand complexes for the hit compounds, 100 ns simulations were performed for each compound. Avicularin-receptor complex (lig-fit-prot) reached equilibrium after the first 30ns, the equilibrium was maintained till the end of the evolution with minimum and maximum values of 1.39 and 2.77Å, respectively. The Cα atoms reached a consistent fluctuation only after 0.5Å and was maintained all through the model. There was no drastic increase in RMSD value, possibly indicating stability of these two systems [42]. Quercetagetin complex, on other hand, showed a steady state from 15ns to 78ns and the oscillation was between 1.6 Å and 3.2 Å. The slight fluctuation of the quercetagetin complex might be due to the number of rotatable bonds of the functional group which protrude outward of the target binding pocket [43].

Interestingly, there was no substantial fluctuation in the loop regions while comparing within models. The amino acid residues in avicularin, hyperoside, sylibin, and harpagoside model oscillated with little fluctuations; 0.5 - 1.5 Å, 0.5 - 1.6 Å, 0.5 - 1.0 Å, and 0.6 – 1.2 Å, respectively. We observed fluctuation in the quercetagetin model with 5.4 Å highest loop. The massive instability of the loops recorded was due to their inherent flexible nature of the ligand, which might be associated with the ligand interactions [44]. The instability of the quercetagetin model amino acid residues corroborates its lig-fit-prot RMSD, with large fluctuation between 76 and 90ns simulation run time. In essence, we found avicularin, hyperoside, sylibin, and harpagoside to be exceptionally stable in the binding active site of *Pf*HT1 with negligible structural orientation and minimum conformational instabilities, rendering them suitable inhibitors.

When we considered the protein-ligand interactions categorized into hydrogen-bonds, hydrophobic, ionic, and water bridges, we observed that all the selected complexes shared a few amino acid residues that were conserved throughout the simulation run time. Moreover, the hit compounds were selective for *Pf*HT1 over human orthologues (hGLUT1). This potentially validates the compounds as drug candidates of interest. Avicularin, sylibin, hyperoside, and harpagoside, among the top five *Pf*HT1 hits, showed excellent polypharmacological possibilities, as evidenced by their docking scores, binding interactions, ADMET characteristics and interactions with receptor site residues.

## Conclusions

Through molecular docking and MD simulation analysis, we have demonstrated that Asn48, Ser315, Ser317, and Val314 are likely essential amino acid residues for *Pf*HT1 inhibition. We also showed that sylibin, hyperoside, harpagoside, and avicularin are stable in the binding site and may efficiently inhibit *Pf*HT1. Furthermore, the ADME properties of the four compounds make them potential druggable molecules which selectively inhibit the *Pf*HT1 receptor over hGLUT1. This opens a new line of investigation for *in vivo* modelling and preclinical assessment of the chemotherapeutic potentials of the identified compounds.

## Acknowledgments

We acknowledge the effort of Olawale J. Ogunleye (Julius-Maximillan University, Wuerzburg, Germany) during the critical examination of the trajectory files and for his contribution to the review of the R scripts.

## Authors’ contributions

AJO and KMO conceived the project. AJO, OAE and KMO carried out molecular dynamics, docking studies and other computational analysis. AJO, FCL and KMO drafted the manuscript. All authors read and approved the final manuscript.

## Availability of data and materials

All data generated or analysed during this study are included in this published article and its supplementary files. The R Script used for data analysis has been deposited in CeGRIB’s GitHub page (https://github.com/CeGRIB/DrugDesign).

## Abbreviations

MMGBSA: Molecular mechanics with generalised born and surface area
*Pf*HT1: *Plasmodium falciparum* hexose transporter 1
hGLUT1: Human glucose transporter
RMSD: Root-mean-square deviation
TIP3P: Transferable intermolecular interaction potential 3 points
RMSF: Root mean square fluctuation
ADME: Absorption, distribution, metabolism, and excretion.
OPLS: Optimized Potentials for Liquid Simulations

## Declarations

### Ethics approval and consent to participate

Not applicable.

### Consent for publication

Not applicable.

## Supporting information

**S1 Fig (A-E). PfHTI protein Root Mean Square Fluctuation (RMSF) with amino acids that participated in protein-ligand contact**

**S2 Fig (A-E). Scatter plot of the atomic displacement parameter (B.factor) against the central carbon atom C-alpha (Cα) showing the type of protein-ligand contacts (bonds) in *Pf*HT1**

**S3 Fig (A-E). Protein Root Mean Square Fluctuation (RMSF) of *Pf*HT1(6m20)**

**S4 Fig. (A-E). Dot and line plot of the Gibb’s Free Energy (wrt) against Protein Data Bank (PDB) residue showing the ligand RMSF of *Pf*HT1(6m20)**

**S1 File. Compounds identified in this study**

